# Decomposing bulk signals to reveal hidden information in processive enzyme reactions: A case study in mRNA translation

**DOI:** 10.1101/2023.05.17.541147

**Authors:** Nadin Haase, Simon Christ, Dag Heinemann, Sophia Rudorf

## Abstract

Processive enzymes, such as polymerases or ribosomes, are often studied in ensemble or bulk experiments through the monitoring of time-dependent signals, such as fluorescence time traces. However, ensemble signals are a superposition of the time traces of all molecules in the reaction and can exhibit less distinct features than individual single-molecule signals due to the stochasticity of biomolecular processes. Here, we demonstrate that under certain conditions, bulk signals from processive reactions can still be decomposed to reveal hidden information about individual reaction steps. Using mRNA translation as a case study for processive biochemical reactions, we explore the limits of least-squares approaches for ensemble fluorescence signal decomposition. Specifically, we show that decomposing a noisy ensemble signal generated by the translation of mRNAs containing more than a few codons represents an ill-posed problem, which can be addressed through Tikhonov regularization. Our findings can help to increase the information content extracted from bulk experiments, thereby expanding the range of these time- and cost-efficient methods.

## Introduction

Many biochemical reactions are driven by processive enzymes that catalyze multiple rounds of reactions while staying attached to the same polymeric substrate molecule. Prominent examples of processive enzymes are DNA and RNA polymerases, ribosomes, cellulases, or endonucleases [1, 2]. To study the kinetics of the biochemical processes driven by processive enzymes, basically two distinct approaches can be taken: single-molecule experiments and ensemble studies [3, 4]. In single-molecule experiments, the action of an individual enzyme is monitored over time, which results in a detailed picture of the step-by-step motion of this molecule (see Fig. 1). Because thermal fluctuations cause differences in the molecular activities, single-molecule experiments have to be performed on a multitude of individual enzymes to obtain a complete picture of the biochemical process. This need for repetition in combination with an often complicated setup can make single-molecule experiments a technical and financial challenge. In comparison, ensemble (bulk) experiments are relatively cost- and time-efficient. Their results have a high statistical power, because they are performed on a large number of enzymes in parallel. However, the signal of an ensemble experiment is typically a superposition of all individual signals, each of which is emitted by a single enzyme in the reaction volume. Due to the stochasticity of processive biochemical processes, these individual signals differ from each other to some extend. Consequently, features or characteristics of the individual signals are blurred or even lost during superposition and, thus, are not present in the ensemble signal [5, 6]. The more time has passed after start of the experiment, the more pronounced is this blurring, even for initially synchronized ensemble reactions. Therefore, ensemble time traces need to be computationally decomposed to reveal hidden information about the underlying enzymatic process [7]. A meaningful computational analysis of ensemble data relies on a mathematical model that fulfills two requirements: First, the model has to reproduce the experimental data and, second, it must do so with an appropriately limited set of parameters. In general, an under-parameterized model lacks the complexity required to reflect the true biochemical process, whereas model over-parameterization leads to overfitting, which causes ambiguous and unreliable results [8, 9]. However, which number of parameters is appropriate is highly case-specific.

**Figure 1:**
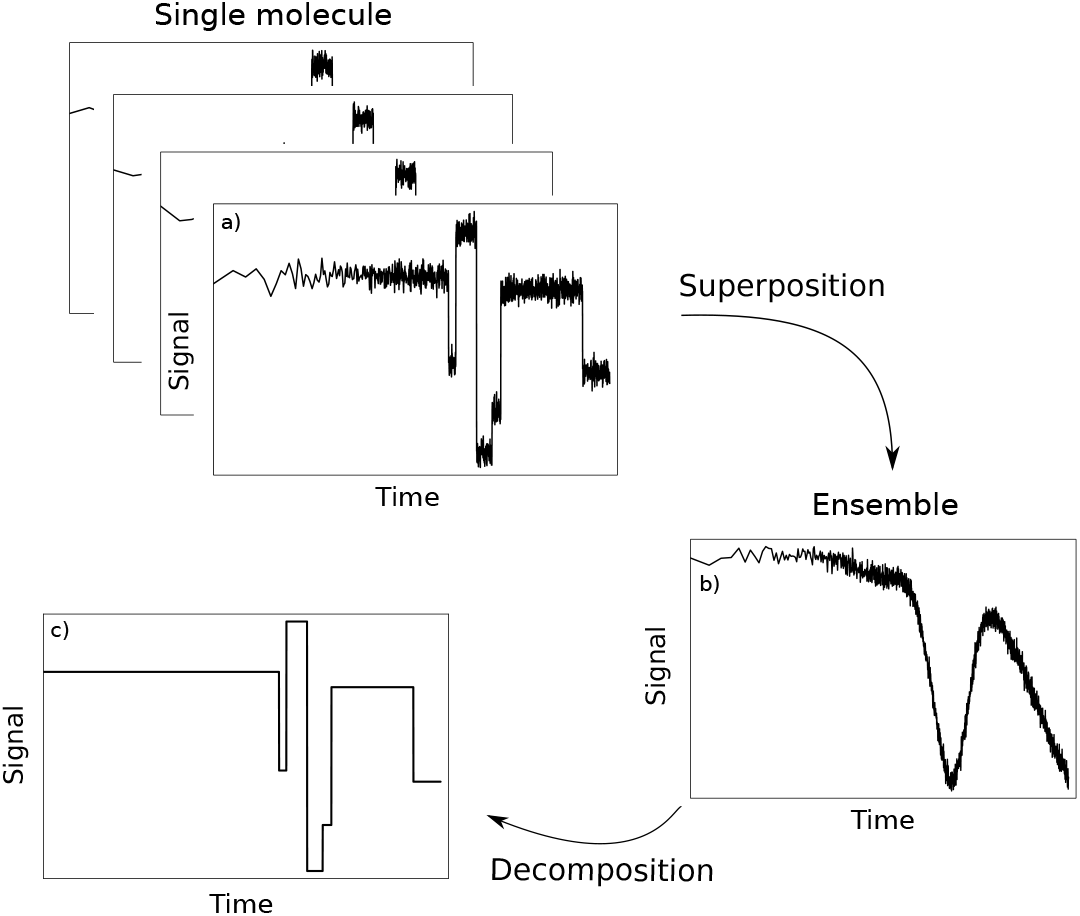
Decomposition of ensemble time traces. a): The action of an individual processive enzyme is monitored over time. Changes in the signal are interpreted to obtain a detailed picture of step-by-step transitions and related kinetics of this molecule. b): Ensemble time trace for a large number of enzymes in parallel. The detected signal is a superposition of all individual signals. Due to the stochasticity of biochemical processes, these individual signals differ from each other to some extend. Consequently, the features of the individual signals are blurred or even lost during superposition. c): The ensemble time trace needs to be computationally decomposed to reveal the hidden information on the transitions of a single molecule.

In the context of molecular biology and biochemistry, fluorophores are used for different purposes, e.g., for visualization of cell compartments or localization of compounds [10, 11]. Because the intensity of the light emitted by a fluorophore depends on the physico-chemical properties of environment *via* quenching and de-quenching, fluorophores can also be applied as molecular probes for state transitions in biochemical reactions [12–17]. For the latter case, it is required that to each state of the reaction process a specific fluorescence intensity can be assigned. We refer to these state-specific fluorescence intensities as “intrinsic fluorescence intensities” (IFIs). The aim of the decomposition of an ensemble fluorescence time trace is to determine these state-specific signal intensities for the individual states. For the sake of clarity, we restrict our analysis to processes that can be assumed to have a fixed (non-stochastic) number of rounds of reactions, i.e., where fluctuations in the processivity can be neglected. Furthermore, we assume that the state-specific signal intensities are the only parameters to be determined by the decomposition procedure and that the kinetic rates related to the action of the enzyme do not need to be inferred from the particular ensemble time trace that is to be decomposed. Of course, determination of IFIs by signal decomposition can be part of an extended experimental and model fitting strategy that in addition also allows for the fitting of kinetic parameters, as we have shown [7].

A well-known example for a processive biochemical process is messenger RNA (mRNA) translation, i.e., the biosynthesis of peptides by molecular machines called ribosomes [18]. Briefly, during translation initiation a ribosome binds to mRNA, i.e., single-stranded RNA which consists of a sequence of trinucleotides called codons that encode for the different amino acids. After binding to the mRNA, the ribosome moves from codon to codon catalyzing the peptide bond formation of the corresponding amino acic molecule and the nascent peptide chain. Translation of the mRNA is finished when the ribosome reaches the stop codon and gets released from the mRNA. Premature drop-off of ribosomes from mRNAs is a rare event (probability of roughly 10^*−*4^ per codon [19]), therefore the processivity of a translating ribosome is mostly simply determined by the length of the mRNA, i.e., the number of codons the mRNA consists of. Except for a few specific cases [20–22], directly assessing the kinetics of mRNA translation in its natural *in-vivo* environment, i.e., in the living cell, is currently infeasible without heavily disturbing the system. However, the movement of ribosomes along mRNAs can be indirectly monitored in *in-vitro* translation experiments by using fluorescent molecules that are attached to the N-termini of the nascent peptides, see ref. [7] for details. The experiments are constructed in a way such that each mRNA is translated only a single time: Once a ribosome reaches the end of the truncated mRNA, it is bound to remain there and does not engage in another round of translation. This yields a unique fluorescence time trace for each translated mRNA that we refer to by the mRNA’s “fluorescence signature”. All ribosomes are initially synchronized and are simultaneosly translating identical mRNA sequences [7]. The ensemble fluorescence time trace (or ensemble fluorescence signature) is a superposition of all fluorescence signals. Translation is a stochastic process, thus the ensemble fluorescence signature obtained from a bulk experiment is much smoother than the individual fluorescence time traces generated by single mRNA translation processes, which consist of discrete gradations [7]. The goal is to find the fluorescence intensities that are associated with the individual states of the translation process from the ensemble fluorescence signature.

In this work, we investigate the feasibility of the decomposition of ensemble time traces that result from the simultaneous action of a multitude of identical, initially synchronized ribosomes. In particular, we consider the typical processivity of the enzymes and analyze its impact on the meaningfulness and truthfulness of ensemble signal decomposition. At first, we show that time traces of low-processivity reactions can be decomposed with good fidelity, in contrast to reactions with higher processivity. We then analyze this phenomenon in terms of the condition number associated to the underlying biomolecular process. Finally, we show that the application of a regularization method can expand the applicability of ensemble signal decomposition to reactions with higher processivity.

## Results

### Decomposition of time traces from low-processivity reactions: Fluorescence signatures of mRNAs consisting of only four codons can be fitted by standard least-squares approach

At first, we study the consecutive translation of only four codons by the ribosome, which can be seen as an enzymatic action with comparably low processivity. To assess the quality of the decomposition results and to precisely quantify the deviations of output (fitted) from input (assigned) parameter values, we use simulated ensemble fluorescence signatures instead of experimentally obtained data: We first gener-ate artificial ensemble fluorescence signatures from mRNAs that consist of four identical codons. Then, we “forget” all IFI values that were used to generate the data and, instead, apply fluorescence signature decomposition to obtain estimates for the IFIs.

For data generation, we simulate translation as a continuous-time Markov process where the states comprising the Markov chain represent the steps that a ribosome undergoes during translation. Each state in the Markov process is associated with a respective IFI. The Markov model used in this work includes 6*n* + 2 states and *n* + 3 assigned IFIs, where *n* is the number of codons of each mRNA. Therefore, the Markov model for translation of an mRNA consisting of four codons includes 26 states and seven IFIs. The IFIs are assigned to the states in a way such that they represent expected changes in fluorescence intensity during the biochemical process of translation, e.g., after translocation of the ribosome to the next codon. Therefore and in the interest of fit parameter reduction some states are assigned the same IFI value. For further information on the Markov model and the assignment of the IFIs, see [7] and Fig. 6 in the Methods. Using our Markov model of translation, we calculate the time-dependent state occupancy probabilities **P**, see Methods. These probabilities express how likely it is to find a ribosome at a specific time in a certain state. Furthermore, we construct IFI input vectors 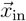 consisting of *n* + 3 random positive real numbers. The simulated input ensemble fluorescence signature 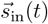 is then given by the sum over all components of the IFI vector 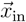 weighted by the time-dependent state occupancy probabilities **P** of the Markov process:

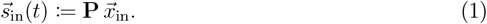

To imitate experiments in which thermal fluctuations modulate the fluorescence signal, we overlay the simulated input fluorescence signature 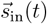 by a Gaussian random noise 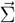 such that resulting final simulated data are given by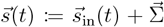, see Fig. 7 in the Methods.

To test our decomposition strategy for the simulated ensemble fluorescence signature, we pretend that all state-specific IFIs are unknown. Analogously to experimental data analysis, where short-term fluctuations are smoothed out, the simulated data 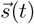 are also smoothed by a moving average operation, i.e., 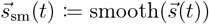 Fig. 2 a) shows an example for a smoothed simulated ensemble fluorescence signature 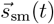.The smoothing step is necessary to fit the model using a least-squares solver to obtain the output IFI vector 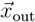

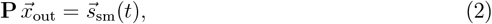

see Methods for details.

**Figure 2:**
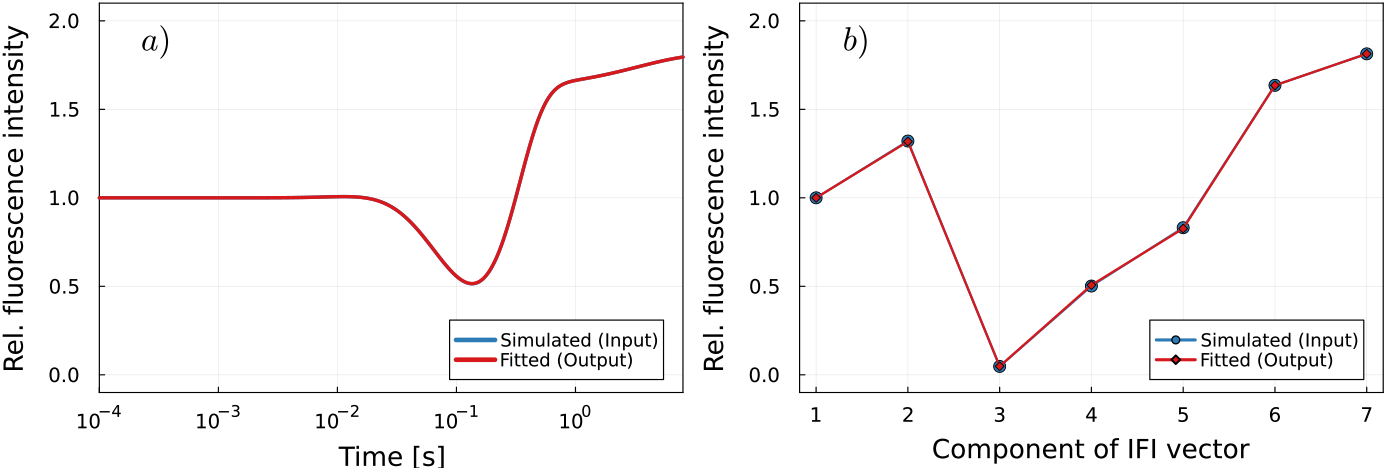
Fluorescence signature of an mRNA that consists of four identical codons for random IFI input vector. a): The simulated fluorescence signature is compared to the best theoretical fit in terms of least squares. The theoretical model and simulated data curves are in perfect agreement. b): Fitted IFIs obtained from the analysis of the fluorescence signature compared to the given IFI input vector.

Ideally, a perfect fit would reveal the IFI values that were used to generate the fluorescence time traces, i.e., 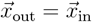. However, due to the added noise, the output and input IFI values 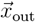 and 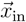 may slightly deviate. Figure S1 in the Supplemental Information shows that the uncertainty is lower in fitted IFI values that correspond to the states in the beginning and towards the end of the studied process. This inhomogeneity in the fit quality is a consequence of the constraints given by the particular “imitated” experimental setup [7]: First, the translation processes are initially synchronized in the sense that all ribosomes simultaneously start to translate the mRNA molecule that they are bound to. By definition, the intrinsic fluorescence intensity that corresponds to the initial state is set to IFI_1_ = 1, i.e., it serves as a reference for the normalization of all fluorescence values. Second, all ribosomes end up and remain in the same last “end” state. Therefore, the IFI corresponding to the “end” state can be well determined as long as the process is monitored long enough, see also Fig. S2 in the Supplemental Information.

In summary, time traces from initially synchronized biochemical reactions with low processivity can be decomposed by standard least-square approaches, and even subtle differences in intrinsic fluorescence intensities of different states can be detected, see Fig. 2 b) and d).

### Standard least-squares approach is not sufficient to decompose time traces from higher-processivity reactions

In this paragraph, we show that signal time traces from reactions with higher processivity generally cannot be decomposed by standard least-square methods and that, instead, more elaborate data analysis methods are required to obtain meaningful fits. To show the limits of the standard least-square approach, we proceed as described in the previous paragraph. However, here we analyze simulated ensemble fluorescence signatures corresponding to translation of mRNAs consisting of 26 identical codons, see Fig. 3 a), instead of just four codons as in the previous paragraph. The Markov model includes 6*n* + 2 = 158 states and *n* + 3 = 29 assigned IFIs, with *n* = 26 the number of codons on the mRNA. Decomposition of these ensemble fluorescence signatures by a standard least-square approach reveals IFI values that greatly deviate from the randomly chosen input IFI values that were used to generate the fluorescence data, see Fig. 3 b). The components of the fitted IFI vector exhibit strong oscillations, span several orders of magnitude, and have negative values. These unreasonable fitting results indicate that the present least-squares problem is ill-posed [23]. To validate this assumption, the singular values *σ*_*i*_ and the condition number *κ*(**P**) of matrix **P** are calculated, see Supplemental Information for definitions and further details: The singular value decomposition of matrix **P** reveals that the singular values *σ*_*i*_ decay gradually to zero, see Fig. S3 in the Supplemental Information. Furthermore, the condition number *κ*(**P**) of matrix **P** is very large, with a value of *κ*(**P**) = 4.8 *×* 10^9^. Both criteria imply that the matrix **P** is ill-conditioned, which means that the computed solution of the ill-posed problem Eq. (2) is potentially very sensitive to perturbations in the input data, i.e., the presence of thermal noise in the simulated ensemble fluorescence signature 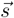 [23, 24]. Fig. 4 a) shows that the condition number grows exponentially with the level of processivity, i.e., the number of repeats of the enzymatic action performed by the processive enzyme. This exponential growth is accelerated when the number of repeats is larger than five. The condition number corresponding to translation of mRNAs consisting of 26 codons is several orders of magnitude higher than the condition number corresponding to translation of four-codons mRNAs. Therefore, solving Eq. (2) for the unknown IFI values in the sense of least-squares is feasible for signals from the translation of 4-codon-mRNAs, but is an ill-posed problem in the case of mRNAs with 26 codons. In addition, the condition number depends on the kinetics of the biochemical reaction, see Fig. 4 b). Here, the kinetics are determined by the elongation rate of the translating ribosomes, which is reflected in the state occupancy probabilities matrix **P**.

**Figure 3:**
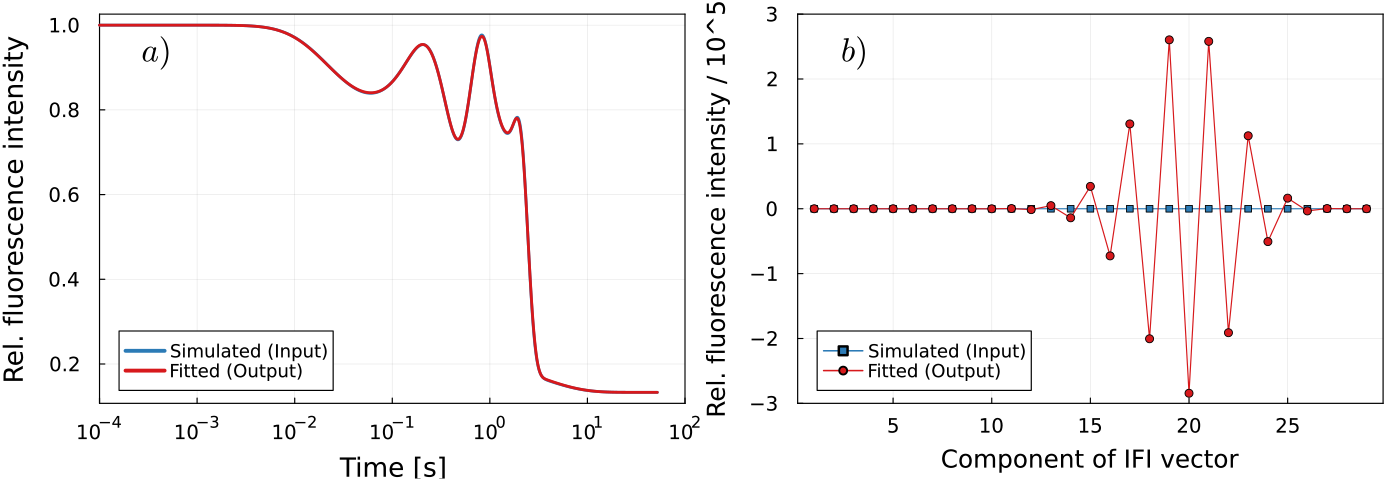
Fitting fluorescence signatures of longer mRNAs reveals signs of an ill-posed problem. a): Fluorescence signature of an mRNA that consists of 26 identical codons for a random IFI input vector. The simulated fluorescence signature is compared to the best theoretical fit in terms of least squares. No regularization is applied. b): Fitted IFIs obtained from the decomposition of the fluorescence signature in a) compared to the given IFI input vectors. No regularization is applied.

**Figure 4:**
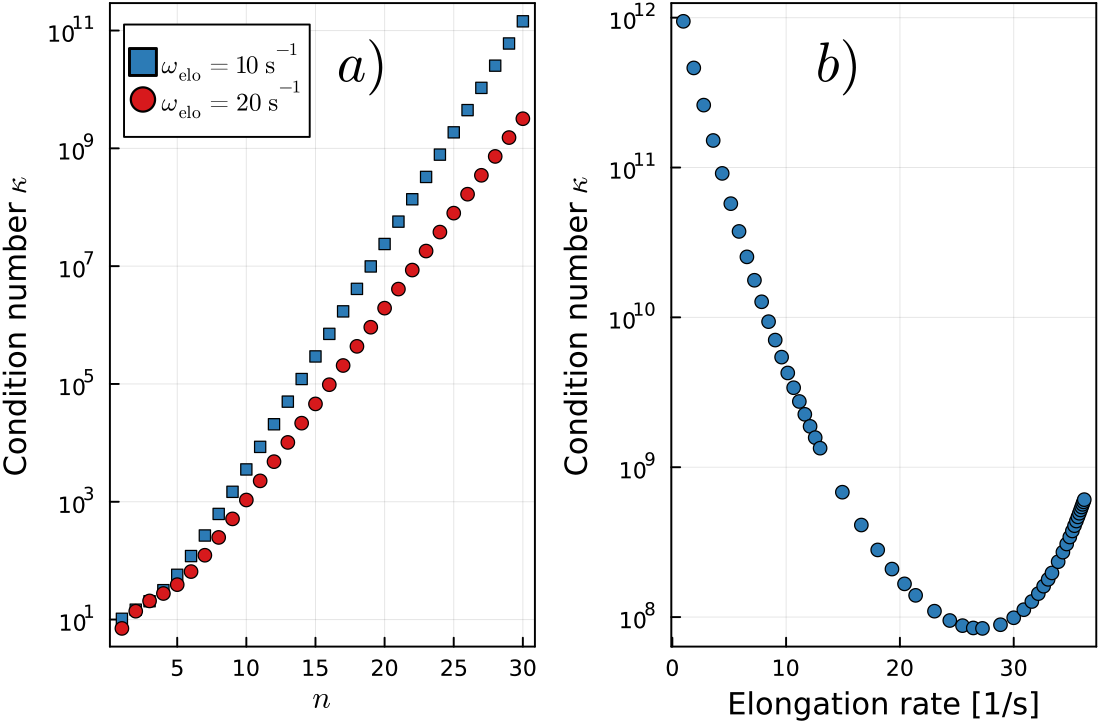
Characteristics of the occupancy probabilities matrix P. a): Condition number *κ* for translation of mRNAs consisting of *n* identical codons with elongation rates *ω*_elo_ = 10 s^−1^ and ω_elo_ = 20 s^−1^. b): Condition number *κ* for translation of an mRNA that consists of 26 identical codons for various elongation rates *ω*_elo_.

### Application of Tikhonov regularization to restore decomposability of signals from high-processivity processes

To find a meaningful solution of a linear ill-posed problem by the method of least squares, the inclusion of additional information is required, which is commonly done by applying Tikhonov regularization [25]. The purpose of regularization is to include side constraints and provide robust methods for choosing the weight given to these side constraints to solve the ill-posed problem, see the Supplemen-tary Information for an introduction to the method. Briefly, a damping is added to filter out the components corresponding to the small singular values *σ*_*i*_ of the matrix **P** [24]. The amount of applied filtering is controlled by the regularization parameter *α*. For larger values of the regularization parameter *α*, stronger regularization is applied, i.e., more filtering is introduced, the fitted solution vector 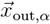 gets smoother and changes only little with higher *α*. Simultaneously, the regularization error increases, which means that some physically meaningful components of the unknown solution are filtered out (regularization error). For a small regularization parameter *α*, the fitted solution vector 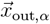 is dominated by terms corresponding to the smallest singular values *σ*_*i*_ and its entries show oscillatory sign changes (perturbation error). In addition, small changes in *α* result in strong changes in the solution 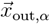. To choose an optimal value for the regularization parameter *α*, the norm of the regularized solution 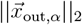 and the residual norm 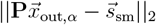 are plotted against each other. For discrete ill-posed problems, the plot in log-log scale shows a characteristic L-shaped curve with the optimal regularization parameter *α* corresponding to the corner of the curve [23, 26]. The optimal regularization parameter *α* balances the regularization and the perturbation error to find a stable solution.

Figure 5 a) shows an example of the L-curve for the decomposition of the fluorescence signature of an mRNA that consists of 26 identical codons. For three different choices of *α*, the corresponding fitted IFI vectors 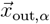 are shown in Fig. 5 b), together with the random input IFI vector 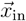. For all choices, Tikhonov regularization led to much more reliable fit results compared to the standard least-square approach (Fig. 3 b)). However, high-frequency features of the input vector are filtered out and thus cannot be recovered by the fitting. For small regularization parameters (see *α* = 0.005 in Figure 5 b)), the perturbation error increases and oscillatory sign changes appear. Nonetheless, the fitted IFI values are still smoother than the input values. Because of this effect, the fit quality depends on the smoothness of the input IFI values. Furthermore, for the same reasons as discussed above for the decomposition of fluorescence signatures of 4-codon-mRNAs, IFI values corresponding to the first and the last states of the translation process are fitted best, see also Fig. S4 in the Supplemental Information.

**Figure 5:**
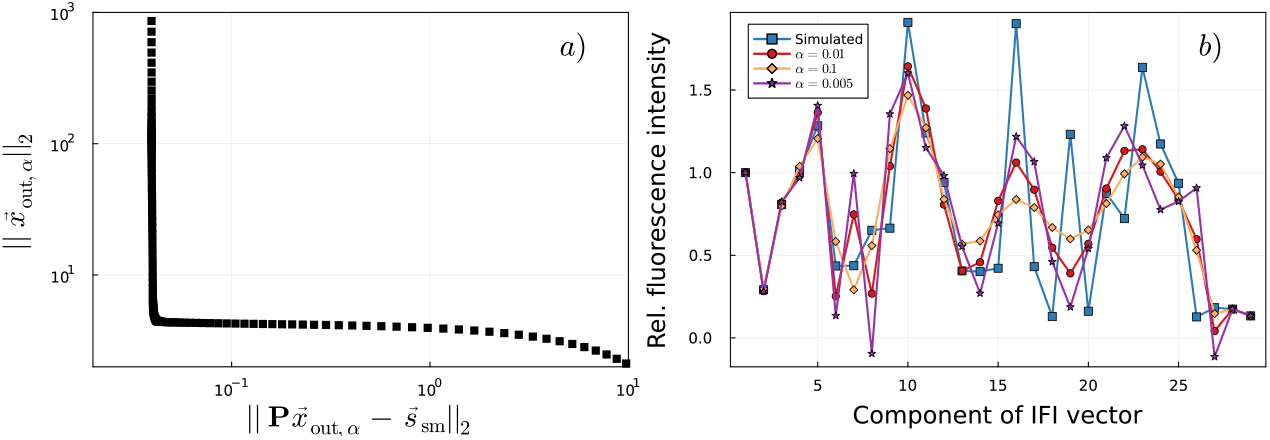
Decomposition of fluorescence signatures of mRNAs consisting of 26 identical codons with Tikhonov regularization. a): For different values of *α*, the norm of the regularized solution 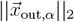 is plotted versus the residual norm 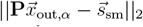. The log-log plot shows the characteristic L-shaped curve. The optimal regularization parameter in terms of trade-off between regularization and perturbation error is *α* = 0.01 and is found by locating the corner of the curve. b): Input and fitted IFI vectors for three different regularization parameter values.

**Figure 6:**
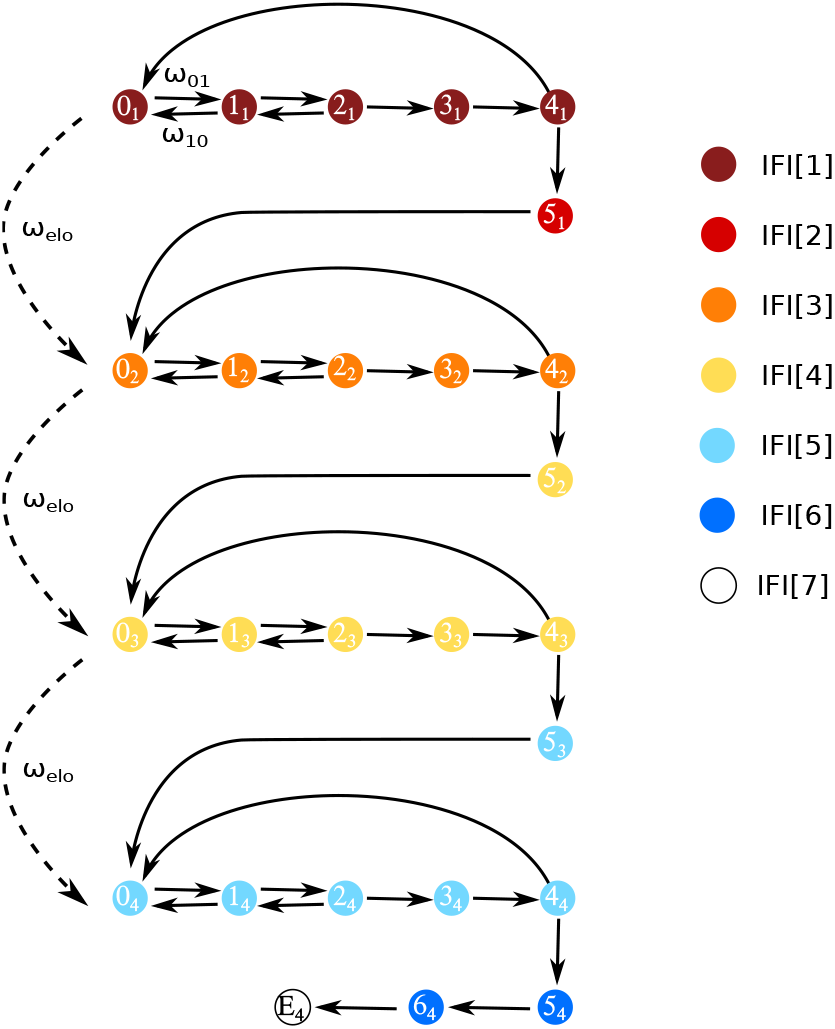
Representation of mRNA translation as a Markov process and assigned IFIs for a mRNA that consists of four identical codons. The Markov model for translation of a short mRNA consisting of *n* = 4 codons includes 6*n* + 2 = 26 states and *n* + 3 = 7 assigned IFIs. The *i*th component of the IFI vector is indicated by IFI[*i*]. Each state in the process corresponds to one biochemically resolved step: State 0_1_ represents the initiation complex with the start codon in the ribosomal A site. States 0_1_-4_1_ describe initial selection, including ternary complex binding and recognition, GT-Pase activation, GTP hydrolysis and rearrangement of EF-Tu. This is followed by tRNA accommodation and peptide bond formation (5_1_). Afterwards, the ribosome translocates to the second codon (0_2_). The elongation cycle is repeated for the next three codons before the ribosome reaches the end of the truncated mRNA (E_4_) where the P site is occupied by the fourth codon while the A site remains empty. See refs. ([29–35]) for more details on the translation process. Possible transitions between the states are indicated by arrows and occur with transitions rates *ω*_*ij*_, which is exemplified by the ternary complex binding rate *ω*_01_ and the unbinding rate *ω*_10_. The average rate of translation for one codon is denoted by *ω*_elo_ and is constant for each of the four codons. For the first codon, changes in the state-specific IFIs are assumed to occur both after peptide bond formation and translocation, and states assigned with identical IFI values have the same color. Note that due to initial synchronization of ribosomes, the two IFI changes on the first codon become visible in the ensemble fluorescence signature. For all following codons only one change in fluorescence intensity after peptide bond formation (after transition from state (4_*k*_) to state (5_*k*_)) is considered to avoid overfitting. The final intrinsic fluorescence intensity E_4_ corresponds to the artificial state of the ribosome at the end of the truncated mRNA. Previously published as the 6*n*2/*n*3 model in [7].

**Figure 7:**
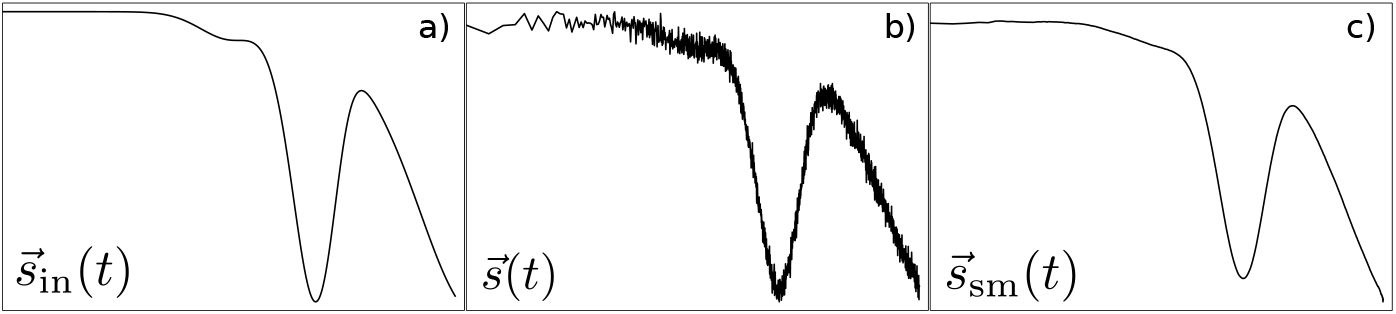
Generation of simulated data. a): Simulated ensemble fluorescence signature 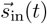 constructed from the random IFI input vector 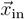 weighted by the time-dependent state occupancy probabilities **P** of the Markov process. b): Superimposition of Gaussian noise to imitate thermal fluctuations. c): The simulated data curve 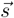 is smoothed using a moving average. Note: To make differences in 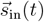 and 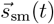 visible, a Gaussian noise with a standard deviation of 0.01 and a moving average with a non-optimal subset size of 200 steps were used in this figure.

## Discussion

We presented a least-squares approach to decompose time-dependent ensemble signals of initially synchronized processive enzymatic reactions and analyzed the limits of the method. The decomposition allows to identify state-specific signal intensities for the individual steps of a reaction, which can be valuable information already by itself or part of a larger model-fitting procedure [7]. To demonstrate the method with a practical example, we modeled mRNA translation as a Markov process and assigned a specific intrinsic fluorescence intensity (IFI) to each state in the Markov chain. We computed the timedependent fractions of ribosomes in the different states of the translation process and determined the corresponding ensemble fluorescence signatures of mRNA translation, including thermal noise. These simulated ensemble fluorescence signatures were then used to test our decomposition method by comparison of input and fitted output IFI values. We found that for reactions with low processivity (corresponding to seven different IFI values in our mRNA translation example), a standard least-squares approach is sufficient to reliably fit IFI values with good precision. In contrast, investigation of model properties in terms of singular values and condition numbers revealed clear signs of ill-posed problems for reactions with higher processivity. However, by using least-square fitting with Tikhonov regularization it is still possible to obtain meaningful solutions. From these observations we draw the conclusion that even before a bulk experiment with initially synchronized processive enzymes is conducted, the following steps can be taken to assess the feasibility of ensemble signal decomposition: First, a model that appropriately describes the stochastic biochemical process of interest needs to be developed, including the assignment of IFIs to the different states of the model. Second, based on this model, the matrix of state occupancy probabilities and its condition number have to be computed (see Eq. (4) and Supplemental Information). For small condition numbers, a standard least-squares approach might be sufficient to decompose ensemble time traces, given that no other factors, such as a low signal-to-noise ratio, prevent a meaningful analysis of the data. Otherwise, regularization is required or, alternatively, strategies must be developed to help decrease the condition number, including, for example, additional measurements to reduce the number of free parameters. Bulk experiments are often much more cost- and time-efficient than methods probing single molecules directly, but they lack the depth of information that the latter approaches achieve. Decomposition of ensemble signals can expand the range of bulk methods by providing a way to extract more detailed information from the same experimental setup.

## Methods

### Modeling of mRNA translation as a Markov process

We modeled mRNA translation as a time-continuous Markov process as is described in ref. [7], [27], and [28]. Most of the known biochemically resolved steps of the translation-elongation cycle are incorporated in the model and the corresponding kinetic rates were investigated by Rodnina and coworkers over the past decades [29–35]. Figure 6 shows as an example the Markov model for translation of an mRNA that consists of four identical codons. By solving the master equations of the Markov process

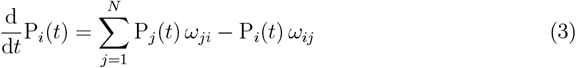

the time evolution of the occupancy probability P_*i*_(*t*), which is the probability to find a ribosome in state *i* = 1, … *N* at time *t*, is obtained for a given set of transition rates *ω*_*ij*_ from state *i* to state *j* (see Table 1). The average rate of translation *ω*_elo_ for one codon can be calculated by first step analysis [36] using the transition rates *ω*_*ij*_ [27]. According to the experimental setup in [7], the translation processes are initially synchronized at state *i* = 0 at time *t* = 0, i.e., the initial condition is

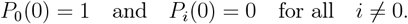

The matrix of occupancy probabilities **P** is defined as

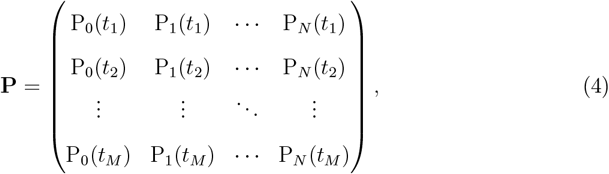

where *N* is the number of states in the Markov process and *M* is the number of considered time points. The time points of the simulation were selected based on the experimental temporal resolution [7].

**Table 1:** Apart from the rates *ω*_45_, *ω*_50_ and *ω*_end_ all transition rates *ω*_*ij*_ are as used in the Markov model description of the translation process in [27]. The average rate of translation *ω*_elo_ for one codon is calculated by first step analysis [27]. The rate *ω*_end_ describes the transition into the artificial end state of the ribosome at the end of the truncated mRNA.

| Rate | Value at 37°C |
| --- | --- |
| $\omega_{01}$ | 350 s<sup>-1</sup> |
| $\omega_{10}$ | 700 s<sup>-1</sup> |
| $\omega_{12}$ | 1500 s<sup>-1</sup> |
| $\omega_{21}$ | 2 s<sup>-1</sup> |
| $\omega_{23}$ | 1500 s<sup>-1</sup> |
| $\omega_{34}$ | 450 s<sup>-1</sup> |
| $\omega_{40}$ | 1 s<sup>-1</sup> |
| $\omega_{45}$ | 13.8 s<sup>-1</sup> |
| $\omega_{50}$ | 53 s<sup>-1</sup> |
| $\omega_{\text{end}}$ | 0.3 s<sup>-1</sup> |
| $\omega_{\text{elo}}$ | 10 s<sup>-1</sup> |

### Generation and fitting of test data

The random IFI input vector 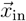 is created using the MATLAB function `rand()` for uniformly distributed random numbers as follows: 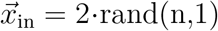, where *n* is the number of required IFI components. The values of the IFI input vector range between 0 and 2, where the first IFI is set to 1. The simulated ensemble fluorescence signature 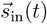 contains *m* = 1… *M* fluorescence values, one for each time point *t*_*m*_. Each individual component of 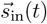 is given by the sum over all components of the IFI input vector 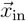 weighted by the time-dependent state occupancy probabilities **P** of the Markov process (see Eqs. (1) and (4)). The simulated fluorescence signature 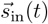 is superimposed with a Gaussian random noise 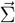 with a standard deviation of 0.005. The influence of the chosen standard deviation on the fitted IFI vector 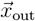 is shown in Fig. S5 in the Supplemental Information. The time points in the simulation and the random noise levels are based on experimental data [7]. The data are smoothed using the MATLAB function `smooth()` with the specified method `moving` and a subset size of 30 data points to obtain 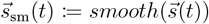. See Fig. 7 for an exemplary representation of 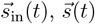. and 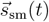 We use the non-negative least-squares solver and `lsqnonneg` in MATLAB to solve the system of linear equations (2) with the components of the IFI vector 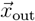 as adjustable parameters.

## Acknowledgments

This work was funded by the Deutsche Forschungsgemeinschaft (DFG, German Research Foundation) – 437345987. Nadin Haase is supported by the Add-on Fellowship of the Joachim Herz Foundation.

Plots were produced using the julia programming language [37] and `Plots.jl` [38].

## Data availability

The MATLAB code for the decomposition algorithm as well as a test data set are avail-able at https://doi.org/10.5281/zenodo.7828502. The following functions were downloaded from the regtools MATLAB package: csvd.m, lcfun.m, l_corner.m, l_curve.m, plot_lc.m and tikhonov.m [39].

