## Supplemental Information for "Decomposing bulk signals to reveal hidden information in processive enzyme reactions: A case study in mRNA translation"

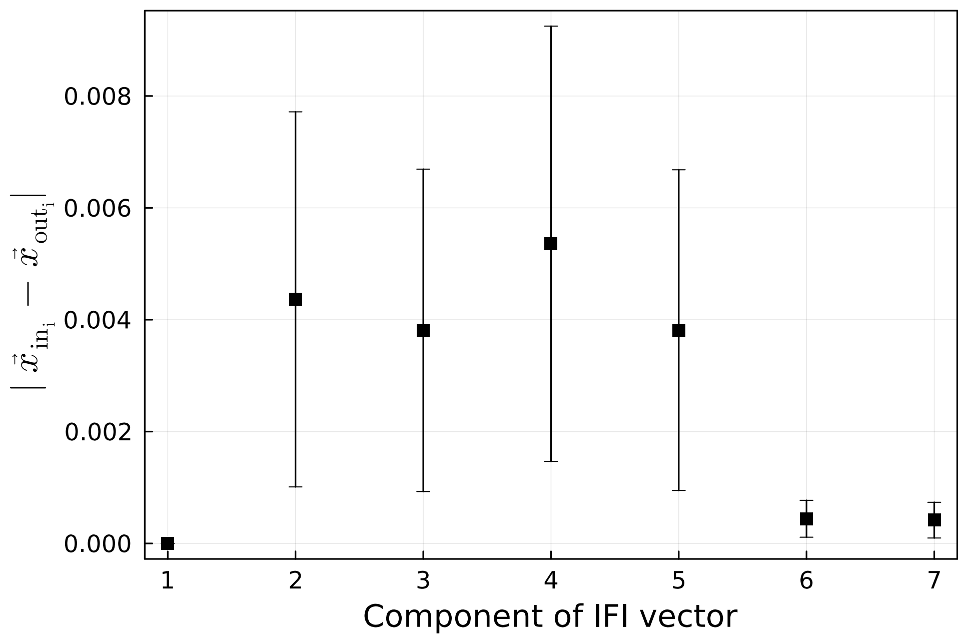


Figure S1: **Component-wise deviation between fitted ($\vec{x}_{\text{out}}$) and given IFI vector ($\vec{x}_{\text{in}}$) obtained from the analysis of the fluorescence signatures of an mRNA that consists of four identical codons.** The shown data is an average for the decomposition of fluorescence signatures simulated from 1000 random input IFI vectors. The first entry of the IFI vector is always set to 1. IFI values related to the last states in Fig. 6 in the main text are fitted with higher reliability.

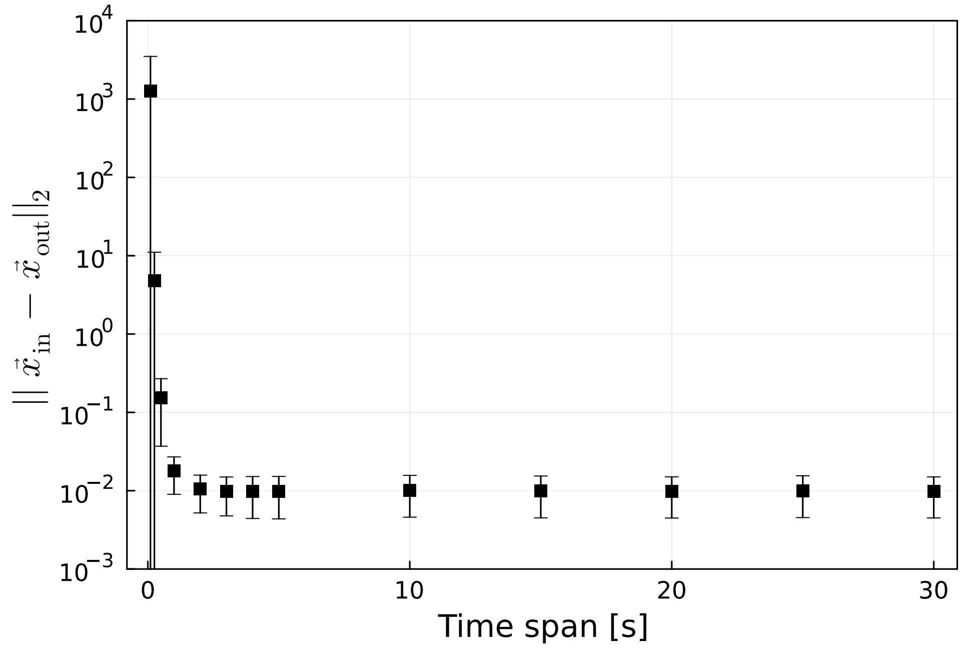


Figure S2: **Deviation between fitted ($\vec{x}_{\text{out}}$) and given IFI vector ($\vec{x}_{\text{in}}$) obtained from the analysis of the fluorescence signatures of an mRNA that consists of four identical codons.** The shown data are averages of the decomposition of fluorescence signatures simulated from 1000 random input IFI vectors. The process was evaluated for different time spans.

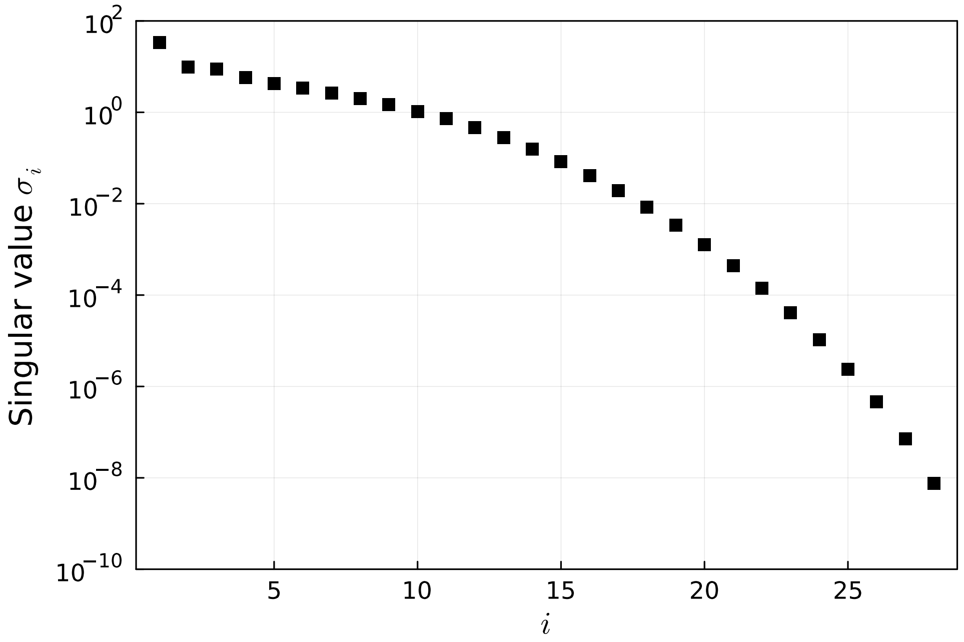


Figure S3: **Singular values $\sigma_i$ of the occupancy probabilities matrix P for translation of an mRNA consisting of 26 codons.** The singular values decay gradually to zero without jumps between consecutive singular values.

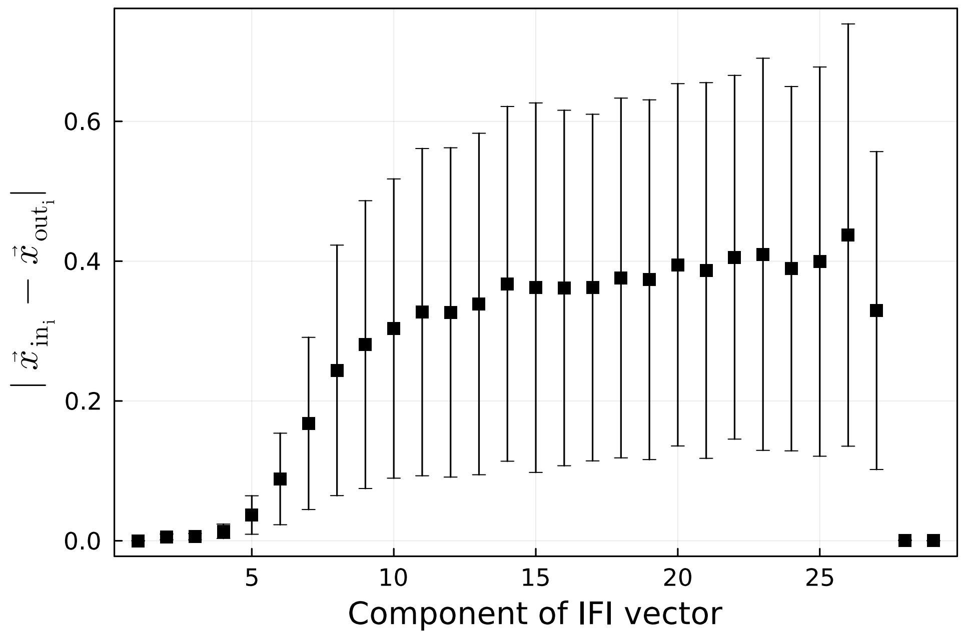


Figure S4: **Component-wise deviation between fitted ($\vec{x}_{\text{out}}$) and given IFI vectors ($\vec{x}_{\text{in}}$) obtained from the analysis of fluorescence signatures of mRNAs that consist of 26 identical codons.** The shown data are averages of the decomposition (with Tikhonov regularization) of fluorescence signatures simulated from 1000 random input IFI vectors. IFI values corresponding to the first and last states of the translation process are fitted with higher accuracy.

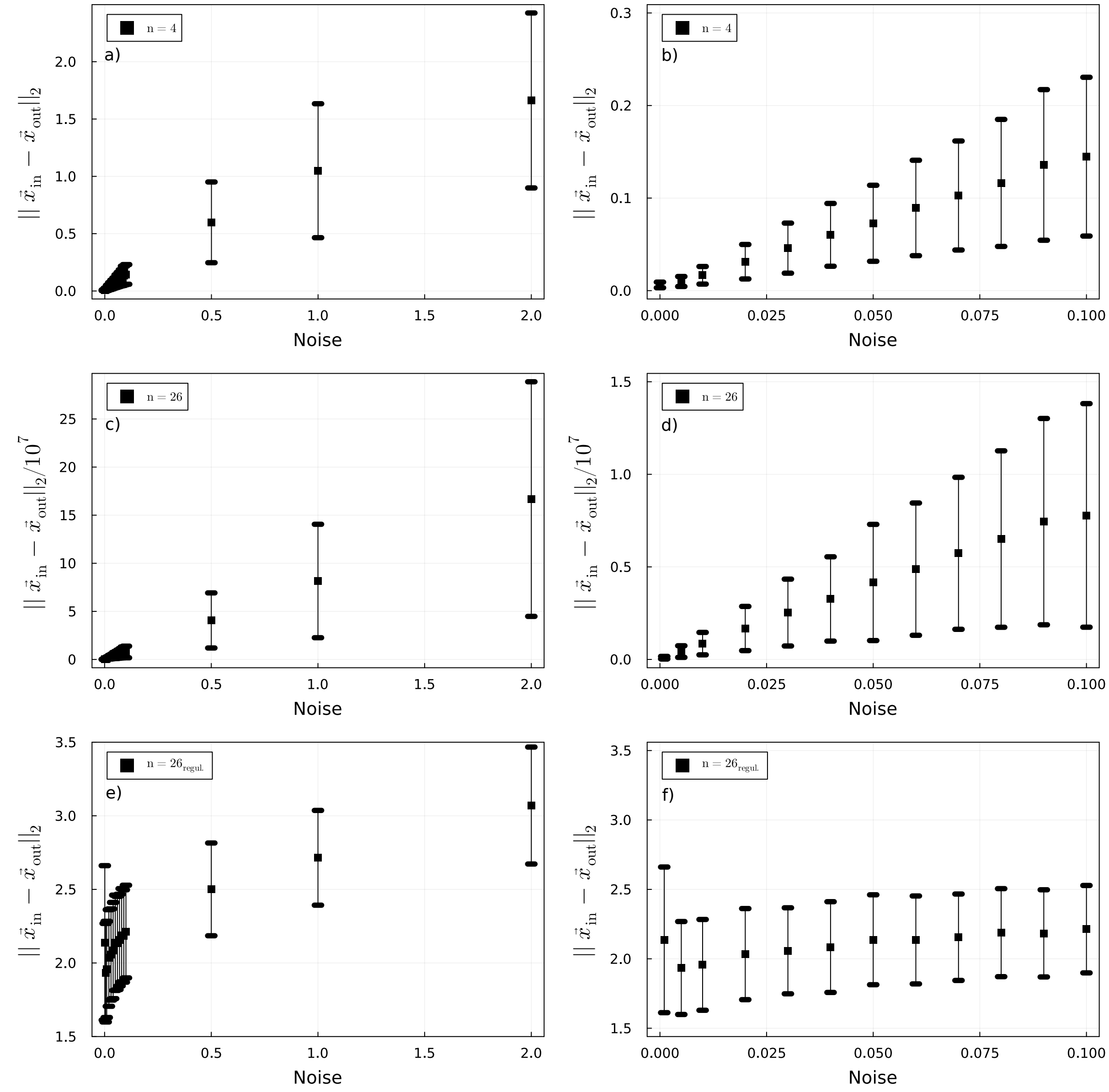


Figure S5: **The deviation between fitted ($\vec{x}_{\text{out}}$) and given IFI vector ($\vec{x}_{\text{in}}$) is plotted versus the standard deviation of the superimposed Gaussian noise $\vec{\Sigma}$.** The data are obtained from the analysis of fluorescence signatures of mRNAs that consist of four identical codons (a and b), of 26 identical codons (c and d) and of 26 identical codons using Tikhonov regularization (e and f). Shown are averages for 1000 random input IFI vectors. Figures on the right side zoom into the range of small standard deviations.

### Singular Value Decomposition and condition number

For a rectangular matrix $\mathbf{A} \in \mathbb{R}^{m \times n}$ with $m \geq n$, the singular value decomposition (SVD) of $\mathbf{A}$ is of the form [1, 2]

$$\mathbf{A} = \mathbf{U}\mathbf{\Sigma}\mathbf{V}^{\mathrm{T}} = \sum_{i=1}^{n} \mathbf{u}_i \sigma_i \mathbf{v}_i^{\mathrm{T}} \tag{1}$$

where $\mathbf{U} = (\mathbf{u}_1, \ldots, \mathbf{u}_n)$ is an $m \times n$ matrix and $\mathbf{V} = (\mathbf{v}_1, \ldots, \mathbf{v}_n)$ is an $n \times n$ matrix with orthonormal columns, so that $\mathbf{U}^{\mathrm{T}}\mathbf{U} = \mathbf{V}^{\mathrm{T}}\mathbf{V} = \mathbf{I_n}$. $\mathbf{\Sigma}$ is an $n \times n$ diagonal matrix with the non-negative singular values of $\mathbf{A}$ of the ordering $\sigma_1 \geq \ldots \geq \sigma_n \geq 0$. The 2-norm *condition number* of $\mathbf{A}$ is defined as [2, 3]

$$\kappa(\mathbf{A}) = ||\mathbf{A}||_2 ||\mathbf{A}^{-1}||_2 = \frac{\sigma_1(\mathbf{A})}{\sigma_n(\mathbf{A})}\,. \tag{2}$$

### Tikhonov Regularization

Consider a system of linear equations $\mathbf{Ax} = \mathbf{b}$ with $\mathbf{A} \in \mathbb{R}^{m \times n}$, $\mathbf{x} \in \mathbb{R}^n$, $\mathbf{b} \in \mathbb{R}^m$, and $m > n$. The vector $\mathbf{x}$ is unknown. Such a problem is called ill-posed if [2, 4]

1. the singular values of $\mathbf{A}$ decay gradually to zero (see Eq. 1), and
2. the condition number of $\mathbf{A}$ is large (see Eq. 2).

The purpose of regularization is to introduce additional information in order to solve an ill-posed problem. In the Tikhonov regularization method a damping is added to filter out the components corresponding to the small singular values [4]. The standard-form version of Tikhonov's method takes the form

$$\mathbf{x}_\alpha = \mathrm{argmin}\{||\mathbf{Ax} - \mathbf{b}||_2^2 + \alpha^2 ||\mathbf{x}||_2^2\}, \tag{3}$$

where $\alpha$ is a positive constant called the regularization parameter. To filter out the contributions corresponding to the small singular values the filter factor $f_i$ is included in the solution [2]

$$\mathbf{x}_\alpha = \sum_{i=1}^{n} f_i \frac{\mathbf{u}_i^{\mathrm{T}}\mathbf{b}}{\sigma_i} \mathbf{v}_i. \tag{4}$$

For Tikhonov regularization, the filter factor is determined by the singular values and the regularization parameter [2]

$$f_i = \frac{\sigma_i^2}{(\sigma_i^2 + \alpha^2)}. \tag{5}$$
